# Peer-Led Team Learning Improves Minority Student Retention in STEM Majors

**DOI:** 10.1101/200071

**Authors:** Jeremy D. Sloane, Julia J. Snyder, Jason R. Wiles

## Abstract

The President’s Council of Advisors on Science and Technology issued a report in 2012 calling for a drastic increase in the number of STEM graduates produced in our country over the following decade if we are to remain economically competitive globally (PCAST, 2012). The report cited the disparity between the diversity among the general public versus that of the STEM professional community and recommended measures to ensure that the women and members of underrepresented racial groups, who together comprise 70% of college graduates but only 45% of college STEM graduates, would become better represented in those fields. This call for action echoed calls by the National Academy of Sciences to expand underrepresented minority participation in STEM at the college level (NAS, 2011). In the following study, we examined whether participation in the Peer-Led Team Learning (PLTL) model in introductory biology influenced the rates of recruitment into STEM and retention in STEM for underrepresented minority (URM) students and for non-URM students. Chi-square analyses reveal that there are significant gaps in STEM recruitment and retention rates between URM and non-URM students, but when these students participate in the PLTL model, no differences in STEM recruitment or retention rates were observed. Additionally, we found that STEM retention rates were significantly improved for URM students who engaged in PLTL.

## Background

In 2012, the President’s Council of Advisors on Science and Technology (PCAST) released a report detailing the need for one million more college STEM (Science, Technology, Engineering and Mathematics) graduates than expected under current assumptions throughout the next decade (PCAST, 2012). The proportion of college graduates that complete a STEM degree has been falling for years, and the proportion of STEM graduates among college graduates is expected to continue to decline. Additionally, the National Academy of Sciences (2011) has identified minority participation in STEM as a national priority, as diversity among participants in STEM fields is necessary to ensure innovation, among other benefits, and to grow a strong and talented science and technology workforce. There is thus a great need to make STEM more accessible to the “underrepresented majority” – the women and members of Underrepresented Minority (URM) groups who constitute 70% of all college graduates but only 45% of STEM graduates (PCAST, 2012).

The first two years of college are critical for STEM persistence. Most students who leave STEM majors do so after taking introductory courses, and, moreover, even high-achieving students often cite uninspiring introductory courses as a reason for switching majors (PCAST, 2012). The PCAST report identifies three main aspects of student experience that affect persistence in STEM: intellectual engagement and achievement, motivation, and identification with a STEM field. It also emphasizes the need to adopt teaching strategies that demand active learning and can improve these facets of students’ experiences with STEM so that the United States can begin to satisfy its own workforce demands.

Peer-led Team Learning (PLTL) is a pedagogical approach that appears to provide much of what PCAST deems necessary to increase student persistence in STEM, including opportunities for intellectual engagement and achievement. Active learning has been documented to improve student learning and reduce failure rates across all STEM disciplines and class sizes (Freeman, Eddy, McDonough, Smith, Okoroafor, Jordt, & Wenderoth, 2014). PLTL is an active learning approach that employs high-achieving undergraduates as peer leaders who facilitate weekly small-group workshops, which the students have the option to attend in addition to or in place of traditional lectures. During PLTL workshops, students work collaboratively on problem sets with their peers and the peer leader. The peer leaders themselves have already taken and been successful in the course and attend weekly training sessions with a learning specialist during which they learn how to facilitate discussions and guide students to their own answers without “teaching” content (Tien, Roth, & Kampmeier, 2002). These workshops promote active learning and engagement on behalf of the students since the students must arrive at the answers to the problem sets themselves. Because PLTL engages students in active learning, active learning has been associated with improved achievement, and achievement in “gatekeeper courses” is closely tied to persistence in STEM, implementing PLTL in an introductory biology course may address intellectual engagement and achievement – the first aspects of student experience that PCAST indicates can affect student persistence in STEM (Alger and Bahi, 2004; Gafney, 2001; PCAST, 2012; Snyder, Carter, & Wiles, 2015). Also, because URM students tend to achieve significantly lower grades in STEM courses than non-URM students and therefore have more potential to gain from active learning approaches, there is reason to believe that URM students may see particular benefits in their STEM retention rates when they participate in PLTL (Rath, Peterfreund, Xenos, Baylis, & Carnal, 2007). We have previously demonstrated that URM students experience particular benefits in their introductory biology grades when they engage in PLTL (Snyder, Sloane, Dunk, & Wiles, 2017).

There is also evidence that instructional strategies that require active learning on behalf of the students can also impact students’ motivation to persist in STEM, the second aspect of student experience discussed by PCAST (2012). Esmaeili and Eydgahi (2014) reported that active learning-based courses have positive impacts on students’ motivation and intention to register for STEM courses. Additionally, providing students with role models in STEM – which the PCAST (2012) report asserts is closely tied to motivation – can influence both recruitment and retention in STEM (Drury, Siy, & Cheryan, 2011). PLTL also provides opportunities for students to interact with peers from similar backgrounds, which has also been associated with motivation to persist in STEM (Ethier & Deaux, 1994). Given that PLTL requires active learning and provides students with role models in the form of peer leaders and opportunities to interact with one another, it may influence student motivation to persist in STEM. Additionally, given that there is a tendency for students to feel isolated and hopeless when not performing well in lecture-based courses, and that URM students tend not to perform as well in STEM courses as non-URM students, PLTL may hold particular benefits for URM students’ motivation to persist in STEM since interacting with peers could potentially alleviate some of those feelings of isolation and hopelessness (NAS, 2011; Swarat, Drane, Smith, Light, & Pinto, 2004).

The third aspect of student experience that the PCAST (2012) report asserts can influence persistence in STEM is identification with a STEM field. Several factors have been documented to influence identification with STEM, including interactions and relationships with peers and faculty, involvement in study groups/discussing and working on course content with peers, and negative racial experiences/degree of feeling included (Anaya, 2001; Chang, Eagan, Lin, & Hurtado, 2011; Espinosa, 2011). The PLTL model provides opportunities for students to work collaboratively with one another on weekly problem sets under the guidance of a peer leader and to feel included in the STEM community, and so may influence each of the above-mentioned factors that are associated with STEM persistence. Additionally, since URM students tend to have difficulty identifying with STEM and since URM faculty are even more underrepresented among peers than URM students are among theirs, PLTL may have particular benefits for STEM identity for URM students (NAS, 2011).

In summary, because PLTL requires active learning, offers role models, and encourages group interactions, it appears to satisfy what the PCAST (2012) deems necessary to increase student persistence in STEM. Moreover, offering PLTL in an introductory course could be an effective intervention at a pivotal point when many students are known to drop out of STEM majors. We predict that PLTL will influence student recruitment into and retention in STEM for students overall, but we also predict that there will be particular benefits for members of URM groups who tend to drop out of STEM majors at higher-than-average rates and may have more trouble identifying with STEM during lecture-based courses (Brown, Henderson, Gray, Donovan, Sullivan, Patterson, & Waggstaff, 2015; Brown, Reveles, & Kelly, 2005).

## Methods

Peer-led Team Learning was offered during the second semester of the Introductory Biology sequence at a large, private university in the American northeast during the. Institutional data for these students was collected three and a half years later, including prior achievement, declared ethnicities, and any declared majors throughout their academic careers. We compared students who participated in PLTL versus those who did not. There was no statistical difference in prior achievement (biology course grades from the previous semester, high school GPA, and SAT) between students who chose to participate in PLTL and those who did not. We considered a STEM major to be any major listed by the National Science Foundation Classification of Instructional Programs for STEM Disciplines (2010). Students were eligible to be “recruited” into STEM only if they did not declare a STEM major upon matriculation to the university and were eligible to be “retained” in STEM only if they ever declared a STEM major. Students were considered “recruited” into STEM if they first declared a STEM major after matriculation. We considered students to have “retained” in STEM if they had remained in a STEM major or had graduated with a degree in a STEM field at the time of data collection—three and a half years after the students completed introductory biology.

Chi-square tests of independence were utilized to examine whether any gaps in STEM recruitment and retention rates existed between URM and non-URM students in the absence of PLTL, whether any differences in these rates were evident if the students participated in the PLTL model, and whether there were any significant improvements in these rates for URM or non-URM students when the students participated in the PLTL model.

## Results

### Recruitment

Table 1 shows the frequencies of URM students, students who engaged in PLTL, and students who were recruited into/retained in STEM majors. Among the students who did not engage in PLTL, URM students were significantly less likely to be recruited into STEM fields than non-URM students (*x*^2^ = 5.415, df = 1, N = 168, p = .020). Among the students who engaged in PLTL, no significant differences in STEM recruitment rates between groups were observed (*x*^2^ = 1.293, df = 1, N = 92, p = .256). There were no significant differences in recruitment rates between URM students who did and did not engage in PLTL (*x*^2^ = 2.126, df = 1, N = 69, p = .145) or non-URM students who did and did not engage in PLTL (*x*^2^ = .895, df = 1, N = 191, p = .344).

**Table 1:** Frequencies of URM students, students who participated in PLTL, students who were recruited into STEM majors, and students who were retained in STEM majors

|  | URM | PLTL | Recruited into STEM | Retained in STEM |
| --- | --- | --- | --- | --- |
| Yes | 88 | 125 | 84 | 114 |
| No | 242 | 233 | 197 | 47 |
| Missing | 28 | 0 | 77 | 197 |

**Table 2:**
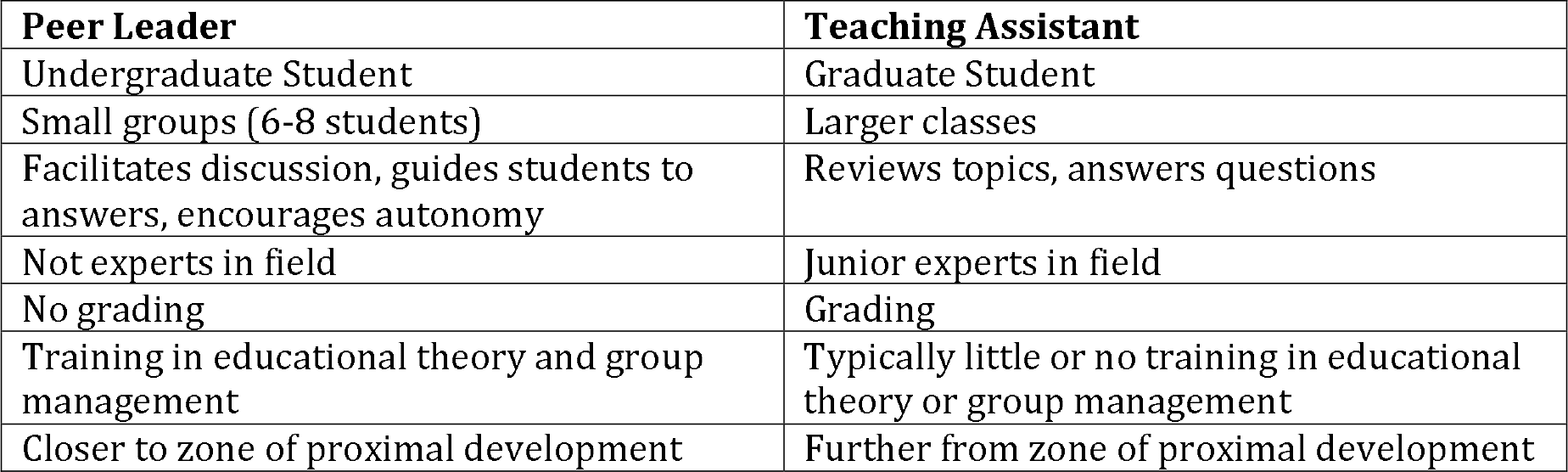
Peer Leader vs. Teaching Assistant

| **Peer Leader** | **Teaching Assistant** |
| --- | --- |
| Undergraduate Student | Graduate Student |
| Small groups (6-8 students) | Larger classes |
| Facilitates discussion, guides students to answers, encourages autonomy | Reviews topics, answers questions |
| Not experts in field | Junior experts in field |
| No grading | Grading |
| Training in educational theory and group management | Typically little or no training in educational theory or group management |
| Closer to zone of proximal development | Further from zone of proximal development |

### Retention

Among the students who did not engage in PLTL, URM students were significantly less likely to retain in STEM fields than non-URM students (*x*^2^ = 6.324, df = 1, N = 95, p = .012). Among the students who engaged in PLTL, no significant differences in STEM retention rates between groups were observed (*x*^2^ = .135, df = 1, N = 53, p = .713) (Figure 1). Additionally, URM students who engaged in PLTL were significantly more likely to retain in STEM majors than those who did not (*x*^2^ = 6.472, df = 1, N = 32, p = .011), while no statistically significant difference in retention rates was observed between non-URM students who did and did not participate in PLTL (*x*^2^ = 3.451, df = 1, N = 116, p = .063).

**Figure 1:**
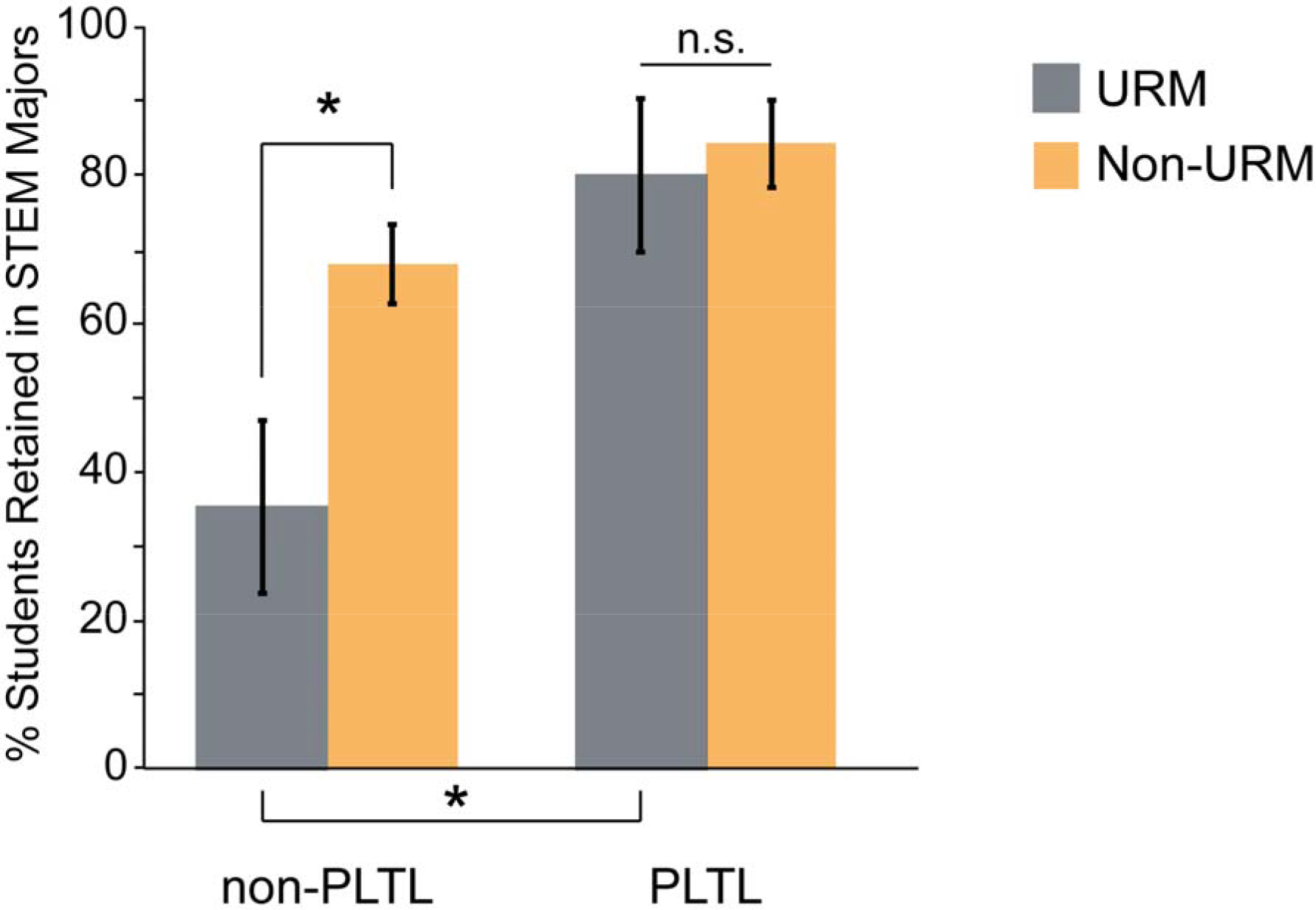
Retention in STEM majors for URM and non-URM students with and without PLTL. Percent of students retained in STEM majors. Values represent percent +/− standard error. Chi-square analyses reveal a significant gap between URM and non-URM students (p = .012) when these students do not participate in PLTL, though no difference in retention rates were observed when students participated in PLTL (p = 0.713)

## Discussion

The results indicate that URM students were significantly less likely to be recruited into or to retain in STEM majors as compared to non-URM students in the absence of PLTL. However, if the students participated in PLTL, no differences in STEM recruitment or retention rates were observed between URM and non-URM students. While there was a significant improvement in STEM retention rates for URM students who participated in PLTL, there was no significant improvement in STEM recruitment rates for these same students.

As a pedagogical approach that demands active learning on behalf of the students, PLTL provides them with a means of making meaning of the course material on their own terms through social interaction with peers. This is associated with better retention of course material and grades in the course (Blake, 2001; Tien et al., 2002). After implementing active learning strategies in a human physiology course, Wilke found that improvements in self-efficacy were associated with increases in course grades for students enrolled in the active learning components of the course (2003). Moreover, URM students have typically earned lower grades than non-URM students in STEM courses (Hunter and Bailee, 2003), and PLTL has been demonstrated to improve grades more for URM students than for non-URM students (Snyder, Sloane, Dunk, & Wiles, 2016). If self-efficacy is tied to student achievement in STEM, student achievement in STEM is associated with student persistence in STEM (as discussed by PCAST), and PLTL increases grades preferentially for URM students in STEM courses, then differences in self-efficacy between URM and non-URM groups may be able to explain the particular benefit of PLTL on URM STEM retention. Future research should attempt to directly measure the effects of PLTL on self-efficacy in association with these other variables to test this hypothesis.

Identification with STEM may also be able to explain why PLTL has particular benefits for retention of members of URM groups. It has been well documented that URM students struggle with identification with STEM, and that this is often a reason that they leave STEM fields (Hurtado, Eagan, & Chang, 2010). Additionally, African-American students who attend Historically Black Colleges and Universities (HBCUs) are far more likely to major in STEM than those at majority institutions (Brown et al., 2015). The PCAST report (2012) indicates that role models are necessary for STEM persistence, and the PLTL model offers role models to students, in the form of peer leaders, who are close to them in age, experience, and identity. In particular, peer leaders are thought to be effective as workshop facilitators and role models because they are considered closer to the students’ “zones of proximal development” and also speak and think more similarly to the students than a typical teaching assistant or professor would (Tien et al., 2002).

There are several limitations of the studies presented in Chapters 2 and 3 that warrant consideration. For students to participate in PLTL, they must attend weekly workshops in addition to the lecture, meaning that they are required to spend more time working on course content. Even though the PLTL workshop materials were made available to the students who did not participate in the model, without having required that the non-PLTL students spend the same amount of time working on course content, it cannot be ruled out that the extra time spent working on the content could be responsible for the differences observed. Additionally, while we attempted to control for selection bias by testing for prior differences in achievement between those who did and did not opt to participate in PLTL, we cannot rule out that the students who opted to participate in PLTL had higher levels of motivation to achieve and persist in STEM than those who did not. Students were awarded extra credit for participating in the model, but this was not included in the grades reported here.

While we are committed to determining which aspects of PLTL may be responsible for the increased STEM retention we have seen among our students, we are no less committed to continuing our use of PLTL in introductory biology if only for the demonstrated benefits toward achievement we have measured among them (Snyder et al., 2016) as well as self-reported attitudes and feelings of confidence we have seen among our peer-leaders. For non-URM students, the PLTL experience at least does no harm in affecting rates of retention in STEM, but for URM students, it appears to make a very significant difference.

## Acknowledgements

The authors would like to thank Beverley Werner for her assistance in the coordination of participant activities and technical skill in data organization and Sarah Hall for assistance with editing figures. We would also like to thank the Biology Department at Syracuse University for their support throughout this project.

